# Discovery of a pentose as a cytosine nucleobase modification in *Shewanella* phage Thanatos-1 genomic DNA mediating enhanced resistance towards host restriction systems

**DOI:** 10.1101/2024.02.27.582347

**Authors:** David Brandt, Anja K. Dörrich, Marcus Persicke, Tabea Leonhard, Markus Haak, Sophia Nölting, Matthias Ruwe, Nicole Schmid, Kai M. Thormann, Jörn Kalinowski

**Affiliations:** Center for Biotechnology (CeBiTec), Bielefeld University, Bielefeld, Germany; Department of Microbiology and Molecular Biology, Justus Liebig University Giessen, Giessen, Germany

## Abstract

Co-evolution of bacterial defense systems and phage counter defense mechanisms has resulted in an intricate biological interplay between bacteriophages and their prey. To evade nuclease-based mechanisms targeting the DNA, various bacteriophages modify their nucleobases, which impedes or even inhibits recognition by endonucleases. We found that *Shewanella* phage Thanatos-1 DNA is insensitive to multiple restriction enzymes and, partially, also to Cas I-Fv and Cas9 cleavage. Furthermore, the phage genome shows strongly impaired basecalling with nanopore sequencing. We characterised the phage adenine methyltransferase TH1_126 in methylase-free *E. coli* ER3413 and derived and confirmed its recognition motif 5’-ATC-3’. Moreover, the data pointed to an additional, much more substantial nucleobase modification. Using LC-MS, we identified a deoxypentose of unknown configuration attached to cytosine as a yet undiscovered phage DNA modification, which is present in Thanatos-1 genomic DNA, likely mediates the observed resistance to restriction endonucleases, as well as a strong reduction in Cas nuclease activity. To elucidate the underlying enzyme functions, we determined structural homologs of Thanatos-1 proteins among known glycosyltransferase folds and experimentally proved a UDP-xylose pyrophosphorylase function of phage protein TH1_063 by *in vitro* enzyme assays.

## INTRODUCTION

Bacteriophages employ a large arsenal of trickery to counter the various bacterial defense systems. It is well known that phages possess modified DNA nucleobases since the discovery of epigenetic modifications in T-even bacteriophages (1). To date, multiple chemical base modifications and the respective enzymatic pathways have been identified and characterized (2). Alternative bases can be identified and analyzed using mass spectrometric methods as LC-MS on digested DNA (2, 3) or, to a limited extent, by direct sequencing of native DNA using third-generation sequencing techniques like SMRT or nanopore sequencing (4–7).

One of the most extensively studied phages, *Escherichia coli* phage T4, regularly adds a hydroxymethyl group to the C-5 of its cytosine residues (hmC), which acts as a protection against bacterial anti-viral defense mechanisms. A common modification pathway in T-even bacteriophages further involves the glycosylation of hydroxymethylated deoxycytosine (dC), where up to two glucose residues are added to hmC by α- and β-glycosyltransferases in the T4 phage (2). This particular covalent DNA modification alone provides complete resistance to *E. coli* restriction-modification (R-M) systems and in addition, the T4 DNA modification also inhibits CRISPR-Cas9 cleavage, which however may depend on the composition of the targeting and tracr RNAs (8, 9). As a second line of defense, *E. coli* T4 possesses a DNA adenine methyltransferase, which methylates GATC sites and also protects the phage DNA from nuclease cleavage (10–12). Recently, another phage epigenetic sugar modification of cytosine, arabinosyl-hmC (ara-hmC), has been identified in *E. coli* phage RB69 (13). The effects on R-M and CRISPR-Cas systems against infection of the phage, however, have not been analyzed yet. This discovery suggests that more DNA base modifications exist that are yet elusive.

Recently, we isolated a lytic phage for the gammaproteobacterium *Shewanella oneidensis* (14). This phage, named *Shewanella* phage Thanatos-1, belongs to the *Tevenvirinae* and shares the typical head-tail morphology. Thanatos-1 has a dsDNA genome of 160.6 kbp encoding about 206 proteins, many of which are of unknown function. An apparent recalcitrance of the phage DNA towards restriction indicated that one or more of the nucleobases may be modified.

Here, we investigated *Shewanella* phage Thanatos-1 (14) with regards to modified DNA nucleobases using nanopore sequencing, as well as LC-MS. We characterized a DNA methyltransferase and identified a so far non-described cytosine modification, which acts as an efficient countermeasure against host nuclease-based defense systems targeting the phage DNA.

## MATERIAL AND METHODS

### Strain cultivation

Bacterial strains used in this study are summarized in Table S1. *Shewanella oneidensis* MR-1 strains were generally cultivated in LB medium at 30 °C or room temperature. *Escherichia coli* strains were grown in LB at 37 °C. For solidification, 1.5 % (w/v) agar was added to the medium. When appropriate, media were supplemented with 50 μg ml^-1^ kanamycin and/or 10 μg ml^-1^ chloramphenicol. Gene expression from plasmids was induced by addition of 0.2 % (w/v) L-arabinose from a 20 % (w/v) stock solution. For the conjugation strain *E. coli* WM3064, media were supplemented with 300 μM of 2,6-diaminopimelic acid (DAP).

### Strain constructions

To determine whether Thanatos-1 is susceptible to CRISPR-Cas-mediated cleavage, we used two different systems, the CRISPR-Cas system IFv from *S. putrefaciens* CN-32 (15) and a system based on Cas9 (16). Corresponding primers and oligonucleotides are listed in Table S2 and S3. To generate an inducible system for the CRISPR IFv system, the corresponding DNA fragment was amplified by PCR from pCas1 (17) with an addition of a C-terminal FLAG tag-encoding sequence to cas6f, the most downstream gene of the cluster, to serve as an expression and production control. The cascade cassette was recombined behind the *araC*-P_BAD_ cassette, which was amplified from pBAD33, by Gibson assembly (18) into EcoRV linearized broad-host range vector pBBR1-MCS5, yielding pCASCADE_RBS_wtCas3. The guide RNA was provided from a second, independent vector. To this end, the appropriate sequence was cloned behind the arabinose-inducible promoter in the vector pBAD33. Both plasmids were verified by sequencing and co-transformed into *S. oneidensis* by electroporation. To generate a Cas9-based system, an appropriate DNA fragment encoding the guiding sequence was cloned into BsaI-linearized vector pTS021 (16) by Gibson assembly. Correct insertion was confirmed by sequencing, and the vector was transformed into *S. oneidensis* MR-1 via electroporation. The oligonucleotides to amplify appropriate spacers from the phage genomes were designed using the CutSPR software (19).

### Phage isolation

To obtain Thanatos-1 and LambdaSo phage particles, 10 ml LB culture of *S. oneidensis* MR-1 ΔLambdaSo ΔMuSo2 was inoculated with 250 μl of an overnight culture and incubated shaking at room temperature until an OD_600_ of 0.4 was reached. After that, 100 μl of phage stock was added and the cultures were further incubated for 24 h. After that, residual cells and cell debris were removed by centrifugation (15,000 × g, 5 min, RT). The supernatant was filtered (0.45 μm), and 1.5-ml-aliquots were stored at 4 °C. The concentration of plaque-forming units was calculated as described below.

### Phage infection assays

To determine the number of infectious, plaque-forming phage particles, 20 ml LB culture of *S. oneidensis* MR-1 ΔLambdaSo ΔMuSo2 was inoculated with 250 μl of an overnight culture and incubated shaking at room temperature until an OD_600_ of 0.4 was reached. If necessary, 0.2 % arabinose was added to induce production of the CRISPR modules in the host strain. Then 15 ml of culture was harvested by centrifugation (10 min, 4000 × g, 4 °C), resuspended in 6 ml LB medium and kept on ice. A serial dilution (10^−1^ – 10^−9^) of the phage suspension was prepared in LB and 100 μl of each dilution was pipetted into a reaction vial. Next 100 μl of the bacterial suspension were added, well mixed with the phage dilution and incubated for 20 min at 30 °C to allow infection of the cells. A control vial did not contain any phages. After incubation, the phage-host culture was mixed with 5 ml of 0.5 % (w/v) soft agar (in LB) and used to overlay a regular LB agar plate. If induction of the plasmid-borne CRISPR system was required, the plates contained kanamycin and/or chloramphenicol and 0.2 % (w/v) arabinose. Plates were incubated at 30 °C overnight. Plaques formed by the phage particles were counted and the amount of PFU per ml preparation was calculated.

As a second assay to determine the number of PFU, spot assays were carried out. To this end, 400 μl of an *S. oneidensis* MR-1 ΔLambdaSo ΔMuSo2 culture were mixed with 7 ml 0.5 % (w/v) LB top agar and spread on regular LB plates (containing kanamycine and arabinose if necessary). After solidification of the top agar, 1.5 μl of a serial phage dilution were carefully spotted onto the surface. Plates were incubated at 30 °C overnight and scored for plaque formation.

### Methylation analysis

For heterologous expression of TH1_126, methylase-free *E. coli* strain ER3413 was obtained from Yale Coli Genetic Stock Center (CGSC#: 14167, New Haven, USA). TH1_126 was amplified from Thanatos-1 DNA by PCR and cloned into pBAD24 vector at the NcoI restriction site using Gibson assembly (18). The Gibson assembly reaction was transformed to *E. coli* DH5α and successful vector construction was verified using Sanger sequencing. Overexpression using ER3413 was induced via overnight cultivation on LB-agar containing 0.1% arabinose. Subsequently, genomic DNA was extracted using Macherey & Nagel Microbial DNA kit (Düren, Germany). Nanopore sequencing was performed on the GridION Mk1 using R9.4.1 flowcells and the Rapid Barcoding kit (RBK-004). Basecalling was carried out using guppy v5.0.11 (available to ONT customers via their community site: https://community.nanoporetech.com/), while tombo v1.5 (20) was used for methylation analysis (https://github.com/nanoporetech/tombo). In a process called “resquiggling”, tombo aligns the raw current signals to the reference sequence. In a second step, tombo’s 6mA *alternative model* method was applied to predict the fraction of modified bases for each adenosine in the genomic reference sequence. By comparing the results of an unmodified negative control (in this case ER3413+pBAD24) to those of the modified dataset, false predictions due to model inaccuracies can be identified. As the exact positioning of the modified base in an interval of roughly five bases cannot be determined due to the inherent nature of nanopore data, the 100 most strongly modified regions of 10 bp were extracted and used as input for MEME motif suite (21). MEME was used with standard settings including “revcomp” mode, which allows a motif to be detected on both strands. MEME output was visualized using WebLogo v3.7.4 (22). The detected, distinct sequence motif (5mATC) was then analyzed by extracting all genomic loci containing NNATCNN and calculating the mean fraction of modified bases across all occurrences of the specific motif, based on the predictions generated by tombo. Additionally, the motif was extended up- and downstream one position at a time, and again, the mean fraction of modified bases for the resulting motifs across all respective genomic loci were calculated and visualized.

### Phage purification and DNA extraction

For chromatographic purification, lysed culture was centrifuged (11,000 × *g*, 10 min, 4°C) and the supernatant was filtered through a 0.2 μm sterile filter. A 1-ml CIMmultus™ OH-1 Advanced Composite Column (pores 6 μM) was used for chromatography on an ÄKTAprime plus system. Phage lysate was diluted 1:1 with 3 M K_2_HPO_4_, KH_2_PO_4_ buffer (pH 7.0) and loaded on the OH column (flow rate 5 ml/min). Buffer A (1.5 M K_2_HPO_4_, KH_2_PO_4_; pH 7.0) was used for washing and for elution a linear gradient from 0 to 100% of Buffer B (20 mM K_2_HPO_4_, KH_2_PO_4_; pH 7.0) was applied. The phage eluate was dialyzed against 10 mM K_2_HPO_4_, KH_2_PO_4_ pH 7.0 buffer using Slide-A-Lyzer Dialysis cassettes (ThermoFisher Scientific, Waltham, USA) for 48 h at 4 °C. Afterwards, phage DNA was extracted using the Phage DNA Isolation kit (Norgen Biotek, Thorold, Canada).

### Exonuclease digestion

Genomic phage DNA was digested using either a combined treatment of DNaseI and ExoI to create single nucleotides or a commercially available nucleoside digestion mix (NEB, Ipswich, USA).

### LC-MS

For LC-MS, the Ultimate 3000 (Thermo Fisher Scientific, Germering, Germany) UPLC system coupled to a microTOF-Q hybrid quadrupole/time-of-flight mass spectrometer (Bruker Daltonics, Bremen, Germany) was used, equipped with an electrospray ionization (ESI) source. For liquid chromatography a SeQuant ZIC-pHILIC 5 μm Polymeric column 150 * 2.1 mm (Merck Millipore, Darmstadt, Germany) was used. For chromatographic separation, 2 μl of the sample was injected and eluent A (20 mM NH_4_HCO_3_, pH 9.3, adjusted with aqueous ammonia solution) and eluent B (acetonitrile) were applied at a flow rate of 0.2 mL min^−1^ by use of following gradient: 0 min B: 90%, 30 min B: 50%, 37.5 min B: 50%, 40.0 min B: 90%, 60 min B: 90%. Mass spectrometry was performed in negative ionization mode. The temperature of the dry gas and the capillary was set to 180 °C. The scan range of the MS was set to 50–1000 m/z. For interpretation of the mass spectrometry data, the software Data analysis 4.0 (Bruker Daltonics, Bremen, Germany) was used.

After dephosphorylation of nucleotides to nucleosides using shrimp alkaline phosphatase (ThermoFisher Scientific, Waltham, USA), LC-MS/MS measurements were performed. In this case, we applied liquid chromatography by using a Bluespher C-18 column 100*2 mm (Knauer, Berlin, Germany). The injection volume was 5 μl and the flow rate was set to 0.4 mL/min. For chromatographic separation, the solutions eluent A (water + 0.1% formic acid) and eluent B (acetonitrile + 0.1% formic acid) was used in the following gradient: 0 min B: 5%, 15 min B: 5%, 20 min B: 80%, 33 min B: 5%. Mass spectrometry was performed in positive ionization mode. The temperature of the dry gas and the capillary was set to 180 °C. The scan range of the MS was set to 50–1000 m/z. MS/MS measurements were performed in MRM mode by first isolating the single mass 360.10 m/z and then applying various collision energies from 10 to 30 eV. For interpretation of the mass spectrometry data, the software Data analysis 4.0 (Bruker Daltonics, Bremen, Germany) was used.

### Heterologous expression and purification of His-tagged Thanatos-1 proteins in *E. coli*

Phage proteins TH1-060 and TH1-063 were amplified from the Thanatos genome and were cloned into the NcoI site of the pBAD24 expression vector. Subsequently, N-terminal 6x His-Tags were added via primer overhangs. Overnight cultures of *E. coli* BL21(DE3)Gold (Agilent, Santa Clara, USA) were inoculated at an OD_600_ of 0.1 into 50 mL LB-Amp in 200 mL shaking flasks at 37 °C with 180 rpm and protein expression was induced at OD_600_ 0.5 with a final concentration of 0.3% L-arabinose, while cultures were cooled to 16 °C and cultivation was continued overnight. Cells were harvested by centrifugation (10 min, 5,500 x g, 4°C), resuspended in Tris-HCl (20 mM, pH 7.5) and lysed on ice using sonication (8×30s). Cell debris was collected at 14,000 g, 4°C and the supernatant was added to Ni-NTA columns and His-tagged proteins were purified according to the Protino Ni-TED protein purification kit (Macherey-Nagel, Düren, Germany).

## RESULTS

### *Shewanella* phage Thanatos-1 dsDNA is resistant to restriction endonucleases as well as Cas9 and Cas-IF cleavage

*Shewanella* phage Thanatos-1 (Thanatos in the following) was isolated using *S. oneidensis* MR-1 as host organism and was hence able to infect and lyse this species (14). *S. oneidensis* possesses a type II R-M which has been shown to decrease the plasmid transformation rate (23), which were obviously not highly efficient towards infection by Thanatos. To determine if *Shewanella* phage Thanatos is generally susceptible to restriction by type II restriction enzymes, we used Thanatos DNA as substrate for various commercially available enzymes and an exemplary assay is shown in Supplementary Figure S1. We found that the Thanatos DNA was resistant to restriction towards a range but not all of type II restriction enzymes.

Another nuclease-based phage defense system that is widely present among bacteria and archaea is the CRISPR-Cascade system (24, 25). Here, the nuclease complex is specifically guided towards potentially invading nucleic acids via short (cr)RNA fragments (26). To determine if Thanatos is susceptible to this phage-defense system, and to potentially set up a genetic system to modify the Thanatos genome *in vitro*, we employed two different types of CRISPR-Cas systems. One was the I-Fv system from the closely related *S. putrefaciens* CN-32 (15), while the second was based on the paradigm Cas9 system from *Streptococcus pyogenes* (27). To this end, the *Sp* I-Fv system comprising the genes *cas3, cas5, cas7* and *cas6f* was placed under control of the arabinose-inducible P_ara_ promoter on a broad-host range plasmid, the corresponding guide RNA was separately expressed from a second plasmid also controlled by the P_BAD_ promoter. For the type II Cas9 system, we adopted a recently described system, which is based on a single plasmid expressing *Cas9* along with the tracr and guide RNAs required for activity (16).

Suitable regions for CRISPR-Cascade interference in phage genomes are defined by so-called protospacer-adjacent motif (PAM), which for both used systems is a ‚GG’ at the 3’-end of the used protospacer. For the *Sp* I-Fv system, two protospacer regions within the Thanatos chromosome were used (within the genes TH1_010, encoding the major capsid protein and the gene TH1_020, encoding a tail lysozyme). As a positive control, we chose two protospacer regions in the genome of *S. oneidenis* phage Lambda (LambdaSo) within the genes SO_2963 (encoding the major capsid protein) and the gene SO_2975 (encoding a protein with unknown function). We previously showed that LambdaSo, which harbors unmodified DNA, is a temperate phage that results in effective lysis in *S. oneidensis* MR-1 strains lacking the prophage and the phage’s integration site (28). The vectors expressing the Cas genes and the gRNA, respectively, were co-transformed into a *S. oneidensis* host strain susceptible to both LambdaSo and Thanatos (*S. oneidensis* ΔLambdaSo ΔMuSo2), which contained no active prophages that may interfere with scoring phage activity by plaque formation. This strain was generally used for all infection experiments. Expression of the respective CRISPR-Cas system was then induced by addition of arabinose. Phage infection efficiency was then determined by soft agar overlay assays applying sequential dilutions of phage particles (Fig. 1). We found that LambdaSo infection and plaque formation was completely abolished in the presence of the CASCADE complex and either of the two LambdaSo protospacers, but not when Thanatos protospacers were used. In contrast, no CRISPR interference was observed when the cells were infected with phage Thanatos, regardless of which protospacer was added, indicating that the I-Fv Cascade system has none or only low efficiency in cutting Thanatos DNA.

**Figure 1:**
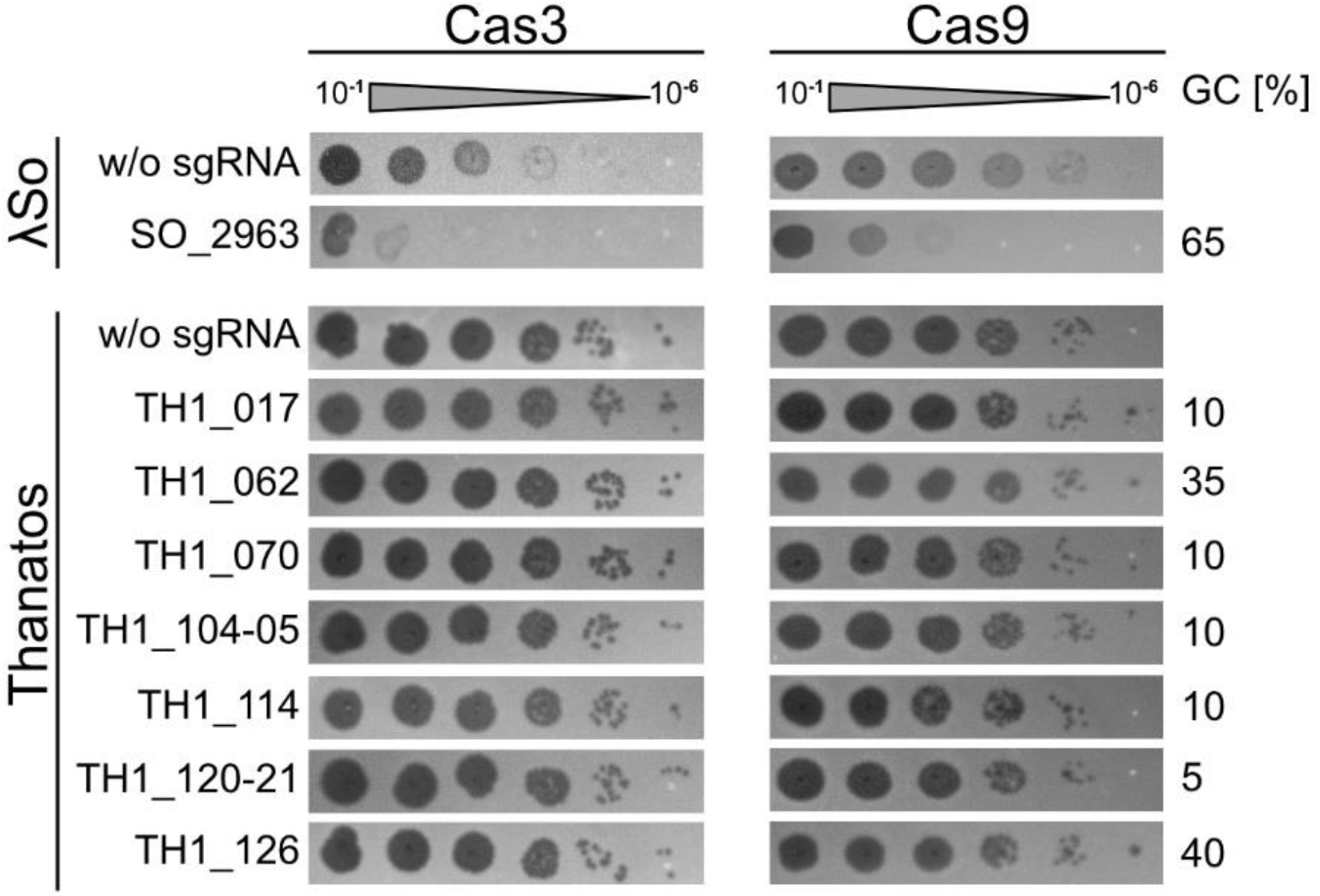
*Shewanella* phage Thanatos exhibits resistance against CRISPR-Cas systems. Shown are spot assays of *S. oneidensis* expressing the CRISPR-Cas I-fv from *S. putrefaciens* CN-32 (Cas3; upper three panels) or Cas9 from *Streptococcus pyogenes* (Cas9; lower three panels). In the left panels, the cells were exposed to increasing dilutions (steps of 1:10) of *Shewanella* phage LambdaSo, and in the right panels to *Shewanella* phage Thanatos accordingly. In each experiment, the cells in the upper panel expressed the corresponding CRISPR-Cas system without any guiding RNA (w/o; negative control). In the lower two panels, a guide RNA targeting the indicated gene was co-produced. The results show that Cas3 from the *S. putrefaciens* IF-v system efficiently decreases lysis by LambdaSo (as a positive control), while no effect occurs on *Shewanella* phage Thanatos. Shown are representative assays of three independent experiments.

Also, the Cas9-based system in *S. oneidensis* showed a significant effect on infection by phage Lambda (Fig. 1). While not as effective as the I-Fv system, occurrence of LambdaSo-derived phage plaques decreased by three to four orders of magnitude, indicating that the Cas9 system is generally active in *S. oneidensis*. However, no effect was observed upon infection by Thanatos, as the number of CFU in assays with Cas9 was indistinguishable from those of the negative controls without guide RNA. Thus, as Cas3 of the I-Fv system, Cas9 is significantly less effective against Thanatos. Taken together, the results show that different bacterial DNA restriction systems do not effectively inhibit Thanatos infection, suggesting that the phage’s DNA is modified in a way that prevents nuclease function.

### Nanopore sequencing of native Thanatos genomic DNA yields low-quality reads

Since Nanopore sequencing has been successfully used for the characterization of DNA modification sites (4, 29), we aimed to employ this technique to elucidate possible DNA modifications of *Shewanella* phage Thanatos. To generate additional data of an unmethylated reference, we used the RPB-004 sequencing kit, which includes PCR amplification prior to sequencing after random transposase fragmentation and adaptation.

Mean quality scores per read clearly differed between native DNA sequencing and PCR-based sequencing (Figure 2) with medians around of 7.5 and 10-12.5, respectively. Converted to percent wrongly called bases, this corresponds to about 20% in the native DNA library and as little as 5% in the PCR-based library.

**Figure 2:**
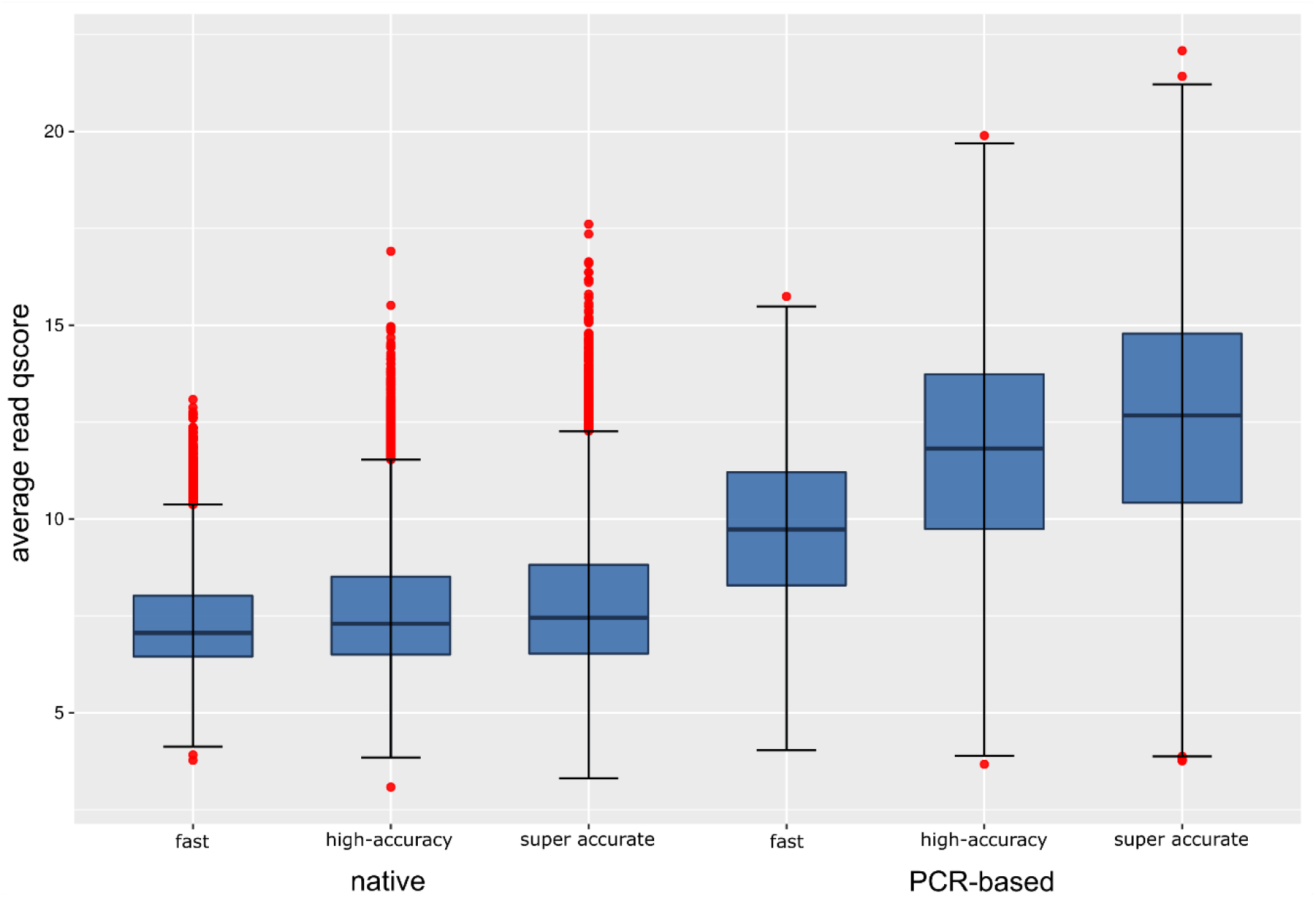
Quality score (qscore) distribution of nanopore sequencing reads as determined by guppy basecaller v5.0.11 using different basecalling modes. Boxplots of average per-read quality scores of sequenced native and PCR-amplified DNAs are shown.

The guppy basecaller provides three basecalling models, employing different algorithms, namely the fast, high-accuracy and super accurate model, which we all used for basecalling. The models differ in achieved raw read accuracy and computation time. Independent of the employed basecalling model, native DNA and PCR-amplified DNA yield sequencing reads of varying quality (Figure 2). Due to the surprisingly low quality of sequenced native DNA and the strong difference to PCR-amplified DNA, we figured that modified DNA may play a role in hindering high-quality nanopore sequencing. We therefore aimed to investigate the genetic repertoire of Thanatos mainly searching for gene products potentially involved in DNA modification.

### Characterization of an adenosine 6mA methyltransferase specifically active on NA*TC sequence motifs

During genome annotation, we discovered a gene (TH1_126) coding for a putative DNA adenine methyltransferase of 273 amino acids. Protein sequence similarity (PSI-BLAST) analysis showed that the most closely related proteins are putative methyltransferase genes from MAGs (metagenome-assembled genomes), classified as *Siphoviridae* sp. viruses with a maximum identity of 39.05 % (95 % query coverage). We aimed to produce this putative phage methyltransferase protein in the methylation-free K-12-derived *E. coli* strain ER3413 (30) to determine a potential methyltransferase activity *in vivo*. TH1_126 was cloned into the arabinose-inducible pBAD vector system (31). Empty vector ER3413 and ER3413 harbouring pBAD::TH1_126 were cultivated on LB agar plates containing 0.1 % (w/v) arabinose. After overnight cultivation and DNA extraction, nanopore sequencing of native DNA was performed. Data analysis was carried out according to a workflow based on tombo (20) as described in the Methods section. Motif detection from 100 sequences (length: 10 bp), which were found to be most strongly modified, yielded a conserved 5’-ATC-3’ motif in 86 sequences (Figure 3). Since MEME was configured to search for a conserved motif on both strands of a given sequence, it reverse-complemented 41 of 86 sequences, which in turn means that 5’-GAT-3’ is also enriched.

**Figure 3:**
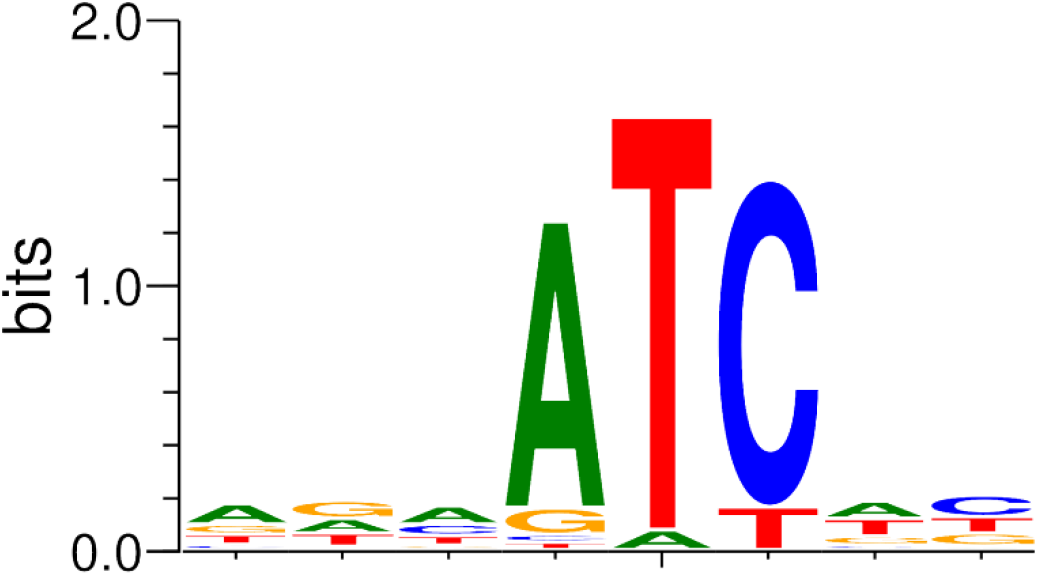
*De novo* detection of modified sequence motifs from Nanopore raw data generated by overexpression of TH1_126 in *E. coli* ER3413. The tool MEME (21) was used to derive a motif from the 100 most strongly modified sequences of length 10. 86 sequences contain the motif “ATC” or the reverse complement “GAT”.

In the following, we will focus on the 5’-ATC-3’ orientation of the detected motif. The control containing only the empty vector plasmid did not yield any conserved motif from the respective sequencing reads in comparison (not shown). We explored up- and downstream extensions of the ATC motif to check for the influence of adjacent bases on the methylation of ATC and found that there likely is no preference of any base upstream of the 5’-ATC-3’ motif (for details see Supplementary Text and Supplementary Figure 2).

The derived methylation site 6mATC overlaps with the recognition site profile of DpnI, which is known to cleave fully methylated GATC sites and to a certain extent also CATC and GATG motifs (32). We did not find any restriction enzyme in REBASE (33) that is known to specifically recognize and methylate a TATC or AATC motif. Therefore, to assess the validity of the bioinformatically derived methylation site, we employed a genomic DNA digestion assay with DpnI, which provided additional proof for the methylation of CATC and GATC motifs (Supplementary Figure 3).

We further conducted a PSI-BLAST search against the *S. oneidensis* MR-1 genome sequence with TH1_126 as input sequence to find possible homologous sequences. We identified SO_0289, which is annotated as Dam family site-specific DNA-(adenine-N6)-methyltransferase, to be a distant homolog of TH1_126 (query coverage 72 %, % identity 24.4 %), which may point to a common evolutionary origin of both phage and host methyltransferases.

### Novel cytosine modification is uncovered by mass spectrometric analyses

Cytosine- or adenosine-specific DNA methylation is not known to impair nanopore sequencing to the extent observed during sequencing of native Thanatos DNA (Figure 2). We therefore hypothesized that there is an additional DNA modification, which would also be consistent with the resistance of Thanatos DNA to restriction endonucleases and reduced Cas cleavage.

To characterize the nature of potential DNA modifications, we employed mass spectrometry. After phage cultivation, we isolated and concentrated the phage fraction using anion-exchange chromatography as detailed in the Methods section. After digestion to single nucleosides, we applied liquid chromatography and mass spectrometry. The chromatogram peaks corresponding to dTMP (321.06 m/z) and dAMP (330.07 m/z) were comparable in peak area, while the dCMP peak was considerably smaller than the dGMP (346.08 m/z) peak. Additionally, a double peak with an m/z of 438.094 was apparent. This unknown peak started to elute after 19.5 min and lasted until 21 min (Figure 4). Its larger mass-to-charge ratio suggested a deoxypentose sugar bound to the cytosine base via an O-glycosidic bond. For further elucidation, we dephosphorylated the nucleotide sample and used tandem MS to fragment the unknown compound in positive ionization mode. After fragmentation, peaks were detected at 128.0450 m/z and 244.0901 m/z. The first peak (128.0450 m/z) most likely corresponds to cytosine with a positively charged oxygen in C-5 position, which possibly stems from a broken O-glycosidic link to a pentose. The second peak (244.0901 m/z) most likely represents the cytosine still bound to the deoxypentose and only devoid of the deoxyribose from the DNA backbone.

**Figure 4:**
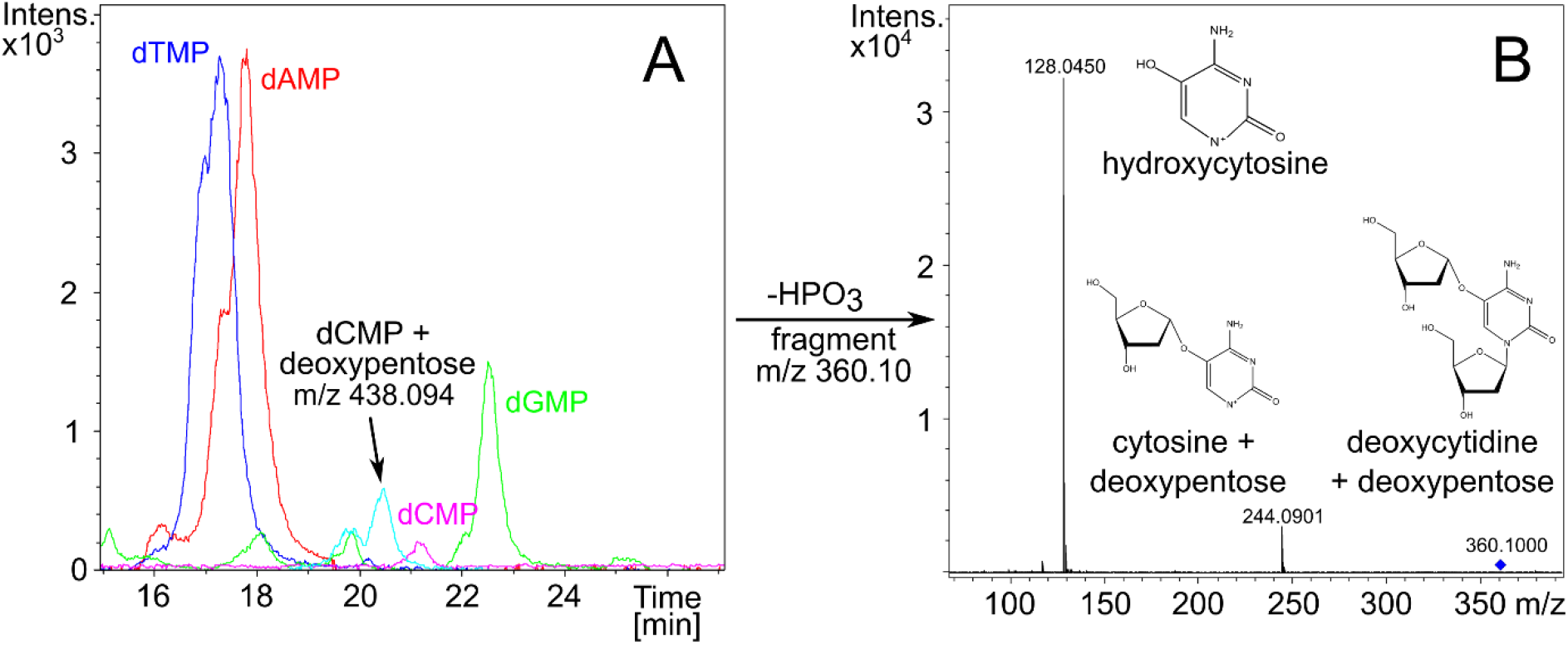
(A) Chromatogram of liquid chromatography analysis with single nucleotides from Thanatos genomic DNA. (B) Mass spectra after isolation and tandem MS fragmentation of m/z 360.10 measured in positive ionization mode after dephosphorylation of nucleotides to nucleosides using alkaline phosphatase.

To date, only one pentose modification of cytosine is known, which is arabinosyl-hmC from Enterobacteria phage RB69 (13). To characterize the pentose configuration of *Shewanella* phage Thanatos, we hydrolyzed the phage nucleosides with trifluoro acetic acid and conducted GC-MS analysis after MSTFA derivatization. As reference standards, we also injected derivatized samples of 2-deoxy xylose and 2-deoxy ribose. We isolated the chromatographic signal of the characteristic pentose mass-to-charge ratio in the range from 306.50 to 307.50 m/z. The chromatogram of the hydrolyzed phage DNA showed a distinct signal containing the above-mentioned m/z. However, the retention time did not match the respective retention times of any of the reference standards. These results lead us to the conclusion that we discovered a novel phage DNA modification, consisting of a deoxypentose bound to the cytosine, which we could not describe further. *Shewanella* phage Thanatos most probably escapes restriction cleavage and Cas attack by attaching the deoxypentose to its cytosine residues.

### Bioinformatic identification of Thanatos candidate genes for cytosine modification

We further aimed to investigate the occurrence of modified nucleotides in Thanatos by searching the genome for putative modification-related gene products. As glycosylation of cytosine residues plays an important role in DNA modification especially in T-even bacteriophages like T4, we focused on gene products possibly involved in this process. Generally, the generation of glycosylated DNA requires activated sugars and a glycosyltransferase enzyme (2). We conducted an analysis of the ORF annotations for cytosine modification-related enzymes and identified three gene products putatively involved in modifying cytosine derivatives (Table 1), TH1_075, TH1_062, and TH1_119.

**Table 1:** Putative nucleotide modification-related genes and gene products in the *Shewanella* phage Thanatos genome identified by protein sequence homology.

| locus tag | Annotated gene function | EC number | Amino acid length |
| --- | --- | --- | --- |
| TH1_075 | dCTP pyrophosphatase | - | 173 |
| TH1_119 | dCMP deaminase | 3.5.4.12 | 174 |
| TH1_062 | Deoxycytidylate 5-hydroxymethyltransferase/thymidylate synthase | - | 237 |
| TH1_060 | Hypothetical protein with glycosyltransferase and phosphotransferase domains | - | 561 |

TH1_075 is predicted to encode a dCTP pyrophosphatase which dephosphorylates dCTP to dCMP. It thus precedes the dCMP deaminase (TH1_119), which catalyzes the formation of dUMP by hydrolytic deamination of dCMP, and, by that, supplies the nucleotide substrate for a thymidylate synthase. TH1_062 is a putative deoxycytidylate hydroxymethyltransferase, which can also be annotated as thymidylate synthase. It may therefore be connected to TH1_119 and TH1_075 by methylating dUMP to generate dTMP, and, in consequence, depleting the host’s dCTP pool.

As our semi-automated genome annotation gave no indication of a glycosyltransferase, we conducted a search for conserved protein domains using CDSEARCH (34) in all annotated protein sequences. By this, we identified hypothetical protein TH1_060 (QJT71744.1; 561 amino acids), which has a predicted glycosyltransferase domain at its N-terminus and a phosphotransferase domain at the C-terminus and may also play a role in DNA modification (Table 1). We further employed structure-based homology search by subjecting the neighboring ORFs of TH1_060, TH1_057 through TH1_065 to ColabFold, an improved implementation of the AlphaFold algorithm (35, 36), figuring that functionally-associated genes likely occur in vicinity of each other. Resulting predicted structures were input into DALI (38) to identify structural homologs in the PDB (39, 40). By this approach, TH1_063 (QJT71747.1; 300 amino acids), was identified as an additional glycosyltransferase candidate enzyme due to its structural homology to known glycosyltransferase enzyme folds (see Supplementary Figure 4 and Supplementary Table 1).

### TH1-063 is a UDP-xylose pyrophosphorylase, possibly also responsible for the generation of activated deoxypentose for cytosine modification

We expressed the two glycosyltransferase structural homologs TH1_060 and TH1_063 with N-terminal 6x His tags in *E. coli* using the arabinose-inducible pBAD expression system (31). After expression and purification, we initially included both proteins in an enzyme assay aiming to assess a putative NTP pentose pyrophosphorylase function, which could provide the activated sugar precursors for cytosine modification. We chose xylose-1-phosphate as a representative phosphorylated pentose precursor, as a phosphorylated deoxypentose was not commercially available to the best of our knowledge. Using LC-MS, we analyzed the reaction mixture consisting of xylose-1-phosphate, one of the five NTPs (ATP, CTP, GTP, UTP, dTTP), pyrophosphatase and either of the phage enzymes TH1_060 and TH1_063. The reaction was incubated at 30 °C for 3 h. In the reaction containing UTP, xylose-1-phosphate and TH1_063, in addition to the chromatogram peaks of the reaction substrates, a peak eluting after 18 min with the prevalent m/z 535.0360 was detected and fragmented using tandem MS in negative ionization mode. Its fragment masses clearly mirror the fragment pattern from tandem MS of UDP-xylose, which was used as a standard (Figure 5). These results show that the TH1_063 protein has a UDP-xylose pyrophosphorylase function, thereby possibly also providing the precursor for the attachment of a deoxypentose to cytosine in a parallel fashion.

**Figure 5:**
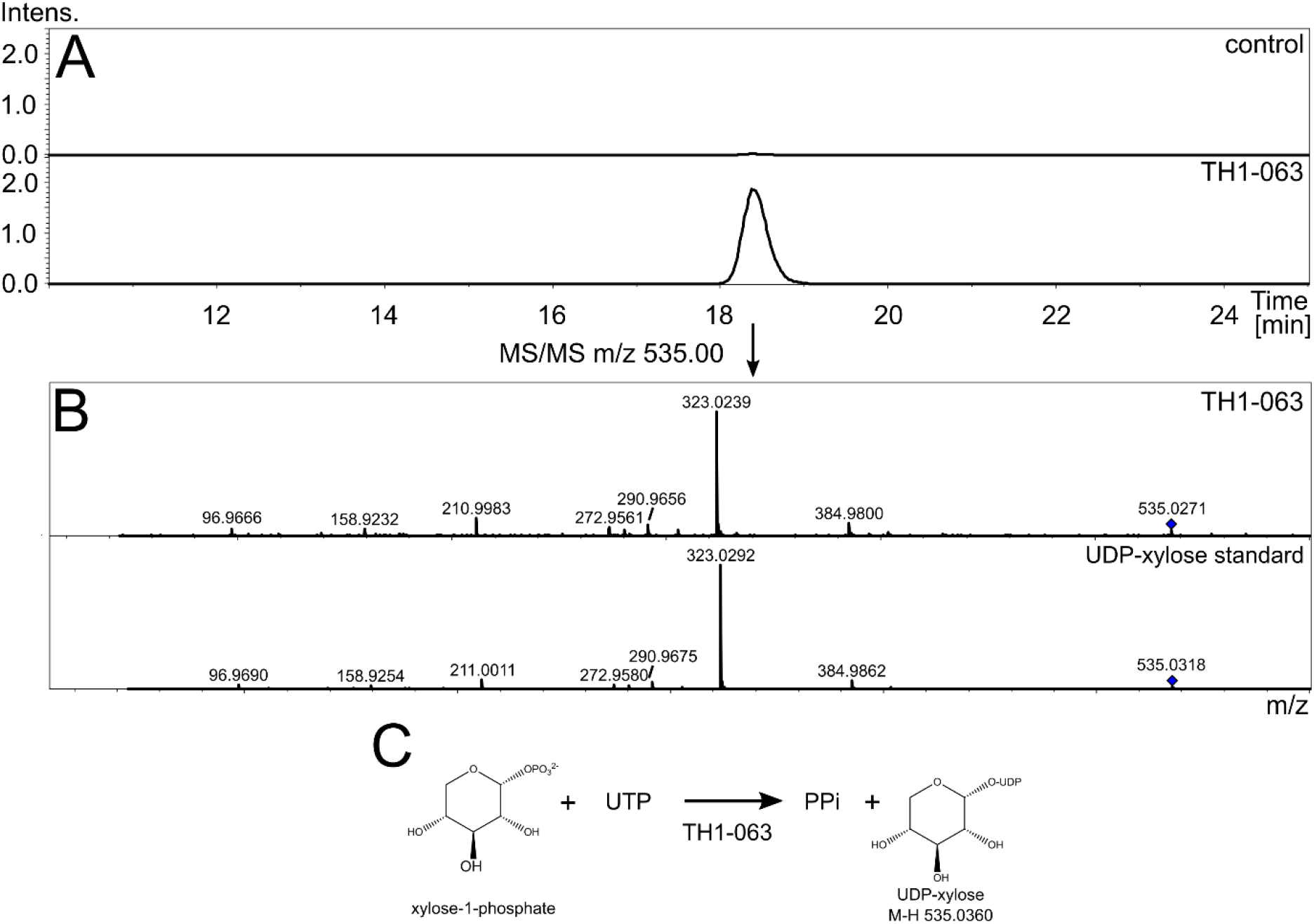
(A) Extracted Ion Chromatogram (negative ionization mode) of m/z 535.0360, which corresponds to UDP-xylose. Upper frame shows no enzyme control, while the assay containing the purified TH1_063 enzyme is shown in the bottom frame. (B) Tandem MS of m/z 535.00 comparing TH1_063 assay and UDP-xylose standard. (C) Reaction schematics of likely UDP-xylose pyrophosphorylase enzyme TH1_063.

Next, we employed both TH1_060 and TH1_063 in an assay containing UDP-xylose and plasmid DNA to examine whether either of the enzymes is able to transfer activated xylose to cytosine. After incubation for 3h at 30°C, plasmid DNA was purified using magnetic beads and digested into single nucleosides at 37°C overnight. Nucleosides were separated and analyzed using LC-MS in positive ionization mode and peaks corresponding to the canonical nucleosides were readily identified. However, we did not detect xylosylated deoxycytosine in our reaction mixture. We further repeated the assays using dCMP, dCTP and cytosine as acceptor substrates but ended up with the same negative results (not shown).

## DISCUSSION

Here, we report the epigenetic characterization of *Shewanella* phage Thanatos including a novel phage adenine methylase and the discovery of a phage DNA modification, which we determined to be a deoxypentose added to deoxycytosine. Previous studies, e.g. on *E. coli* phage T4, showed that nucleotide modifications at the cytosine can effectively decrease the efficiency of CRISPR-Cas systems and thereby protect the phage DNA within the critical time period after injection into the host cell (9, 41– 43). Our results strongly indicate that this is similarly true for the deoxypentose modification of deoxycytosine of *Shewanella* phage Thanatos, at least against the tested restriction by R-M systems and interference by Cas9 and a I-Fv system.

When using nanopore single-molecule sequencing for modification detection, we noticed that high-quality basecalling of sequencing data from Thanatos was only possible when PCR-amplifying the genomic DNA prior to sequencing. Read sequences determined from native DNA were of poor quality and impossible to align to the Thanatos genome sequence, which made modification detection from nanopore raw data unachievable. Interestingly, signal-level data yielded the impression of long reads, while the guppy basecaller only provided short basecalled sequences. This means that pentosyl-deoxycytosine residues pass through the nanopore during sequencing and induce measurable voltage changes, but basecalling models are not equipped to call the respective base sequence. Strongly reduced read quality to the extent of complete inability to map the sequencing reads has also been described recently with nanopore sequencing of phage RB69 DNA, which contains arabinosyl-hmC (44). This challenge remains open for research efforts since the training of basecalling models for bases that strongly differ from the four consensus DNA bases is not easily possible to date, because nearly all tools for modified base detection rely on reference alignment after basecalling (5).

Initially, we had identified the protein TH1_126, which was predicted to be an adenine methylase, and characterized the enzyme by production in the methylase-free *E. coli* strain ER3413 and subsequent nanopore sequencing of the strain’s genomic DNA. Methylation calling and motif analysis yielded a 5’-ATC-3’ recognition site, which was confirmed using a DpnI restriction assay. Since DpnI only recognizes GATC and CATC motifs, we would have preferred to have further proof of AATC and TATC (or even directly for ATC) methylation. However, there were no suitable restriction enzymes which we could use for the respective assay. *Shewanella oneidensis* MR-1 possesses two regularly methylated palindromic motifs (G<u>A</u>TC and ATCG<u>A</u>T) (45) which both contain the sequence motif targeted by the phage methylase. This could point to a protective effect of methylation from host restriction and to a common evolutionary origin of host and phage methyltransferases, which is also supported by the homology of TH1_126 and host methyltransferase SO_0289, which targets GATC motifs (45). However, since *S. oneidensis* MR-1 does not contain any known restriction nuclease cleaving unmethylated GATC motifs (45), methylation by TH1_126 may rather protect the Thanatos genome from cleavage when infection takes place in another host organism. As DNA methylation in *S. oneidensis* MR-1 also plays a role in processes like DNA mismatch repair and regulation of genome replication (45), methylating its genomic DNA may also help Thanatos in synchronizing itself with the host.

LC-MS analyses did show detectable amounts of mA in digested Thanatos DNA, implying that phage methyltransferase TH1_126 was active, at least to a certain extent. It may as well serve another function during phage infection e.g. methylation of host DNA. It is known from the T4 phage that mutants devoid of glucosyl-hmC have a higher methylation frequency of 5’-GATC-3’ motifs, suggesting an inhibitive effect of glucosyl-hmC on adenine methylase (9), which may also be the case with the Thanatos deoxypentose cytosine modification.

Because methylated DNA bases are generally not known to dramatically impair nanopore sequencing, we searched for additional DNA modifications. We found that the Thanatos genome encodes putative dCMP deaminase, dCTP pyrophosphatase and deoxycytidylate hmC transferase/thymidylate synthase enzymes, which are likely involved in nucleotide metabolism or modification. The presence of these three enzymes pointed at a potential modification at the cytosine nucleotides. Accordingly, by DNA digestion and single nucleoside analysis using mass spectrometry techniques, we confirmed such a cytosine modification, which we determined to be a deoxypentose moiety. Our data from LC-MS clearly shows that the deoxypentose is bound directly to the cytosine base via an O-glycosidic bond. This contrasts the findings from T-even bacteriophages T2, T4 and T6, where glucose is bound via an additional hydroxymethyl group to the nucleobase (9, 46).

While three phage-encoded proteins, TH1_062, TH1_075 and TH1_119, are good candidates to be involved in the synthesis of such a modification, candidate enzymes for the generation of activated sugar and a sugar transferase were lacking. Using CDSEARCH, we additionally identified hypothetical protein TH1_060, with both predicted glycosyltransferase and phosphotransferase domains. BLASTP analysis returned 106 putative sequence homologs (cut-offs: 70% query coverage, 40% identity) from different T4-like phage genomes with the best hit being a hypothetical protein from *Yersinia* phage vB_YenM_TG1 at 55.81% identity (YP_009200315.1). The corresponding gene products were classified as hypothetical proteins, “bifunctional protein”, or “NTP-transferase domain-containing protein”. None of these proteins has been characterized experimentally. However, the related *Escherichia* phage RB69 possesses coding region ORF052_053c, which is a homolog of TH1_060, with an amino acid identity of 53.2 % with 100 % query coverage between ORF053_052c and TH1_060. In *Escherichia* phage RB69, ORF052_053c is presumed to play a role in generation of UDP-arabinose, which is essential for the formation of arabinosylated cytosine (13), which also hinted at a possible role of TH1_060 in the generation of activated xylose necessary for subsequent sugar transfer to the cytosine.

The AlphaFold software (35, 36) was used to predict protein structures of proteins TH1_057 through TH1_065. Both TH1_060 and TH1_063 structurally resembled known nucleotidyltransferase and glycosyltransferase proteins and were therefore expressed in *E. coli* and characterized using *in vitro* enzyme assays. We could show that TH1_063 serves as a UDP-xylose pyrophosphorylase, thereby demonstrating the predictive power of AlphaFold (36), which allows protein homology comparisons beyond the sequence alone. After proving the sugar-activating function of TH1_063, one can also speculate that it would also accept a deoxypentose like deoxyribose-1-phosphate as a substrate. This putative activity will be subject of further studies.

We could not identify the function of TH1_060 *in vitro*, which may be a bifunctional enzyme possibly involved in transferring the deoxypentose to the cytosine. Another issue is posed by the hydroxyl group we found at the cytosine after tandem MS fragmentation, as we currently have no valid hypothesis on how the hydroxyl group is attached to the modified cytosine. Therefore, further studies are required to fully identify the protein machinery, which synthesizes the modified cytosine and its precursors and may also include host enzymes, as demonstrated for the *Escherichia* T4 phage (41, 47, 48).

In summary, our study expands the array of general and phage-mediated DNA modifications and demonstrates that the full gamut of DNA modification-based phage-defense systems is still elusive. It remains to be shown whether a direct O-linked deoxypentose modification of cytosine is more frequently found in phages and what biological benefits are associated with this modification. Such studies are currently under way.

## ACKNOWLEDGEMENT

We thank Marina Simunovic for critical reading and correction of the draft manuscript.

## REFERENCES

1. Luria, S.E. and Human, M.L. (1952) A nonhereditary, host-induced variation of bacterial viruses. J. Bacteriol., 64, 557–569.

2. Weigele, P. and Raleigh, E.A. (2016) Biosynthesis and Function of Modified Bases in Bacteria and Their Viruses. Chem. Rev., 116, 12655–12687.

3. Lee, Y.J. and Weigele, P.R. (2021) Detection of Modified Bases in Bacteriophage Genomic DNA. In Methods in Molecular Biology. Humana, New York, NY, Vol. 2198, pp. 53–66.

4. Tourancheau, A., Mead, E.A., Zhang, X.S. and Fang, G. (2021) Discovering multiple types of DNA methylation from bacteria and microbiome using nanopore sequencing. Nat. Methods, 18, 491–498.

5. Xu, L. and Seki, M. (2020) Recent advances in the detection of base modifications using the Nanopore sequencer. J. Hum. Genet., 65, 25–33.

6. Wallace, E.V.B., Stoddart, D., Heron, A.J., Mikhailova, E., Maglia, G., Donohoe, T.J. and Bayley, H. (2010) Identification of epigenetic DNA modifications with a protein nanopore. Chem. Commun., 46, 8195–8197.

7. Rand, A.C., Jain, M., Eizenga, J.M., Musselman-Brown, A., Olsen, H.E., Akeson, M. and Paten, B. (2017) Mapping DNA methylation with high-throughput nanopore sequencing. Nat. Methods, 14, 411–413.

8. Yaung, S.J., Esvelt, K.M. and Church, G.M. (2014) CRISPR/Cas9-mediated phage resistance is not impeded by the DNA modifications of phage T4. PLoS One, 9, e98811.

9. Bryson, A.L., Hwang, Y., Sherrill-Mix, S., Wu, G.D., Lewis, J.D., Black, L., Clark, T.A. and Bushman, F.D. (2015) Covalent modification of bacteriophage T4 DNA inhibits CRISPRCas9. MBio, 6.

10. Schlagman, S.L. and Hattman, S. (1983) Molecular cloning of a functional dam+ gene coding for phage T4 DNA adenine methylase. Gene, 22, 139–156.

11. Yang, Z., Horton, J.R., Zhou, L., Zhang, X.J., Dong, A., Zhang, X., Schlagman, S.L., Kossykh, V., Hattman, S. and Cheng, X. (2003) Structure of the bacteriophage T4 DNA adenine methyltransferase. Nat. Struct. Biol., 10, 849–855.

12. Zinoviev, V. V., Evdokimov, A.A., Gorbunov, Y.A., Malygin, E.G., Kossykh, V.G. and Hattman, S. (1998) Phage T4 DNA [N6-Adenine] Methyltransferase: Kinetic Studies Using Oligonucleotides Containing Native or Modified Recognition Sites. Biol. Chem. Hoppe. Seyler., 379, 481–488.

13. Thomas, J.A., Orwenyo, J., Wang, L.X. and Black, L.W. (2018) The odd “RB” phage—identification of arabinosylation as a new epigenetic modification of DNA in T4-like phage RB69. Viruses, 10.3390/v10060313.

14. Kreienbaum, M., Dörrich, A.K., Brandt, D., Schmid, N.E., Leonhard, T., Hager, F., Brenzinger, S., Hahn, J., Glatter, T., Ruwe, M., et al. (2020) Isolation and Characterization of Shewanella Phage Thanatos Infecting and Lysing Shewanella oneidensis and Promoting Nascent Biofilm Formation. Front. Microbiol., 11.

15. Dwarakanath, S., Brenzinger, S., Gleditzsch, D., Plagens, A., Klingl, A., Thormann, K. and Randau, L. (2015) Interference activity of a minimal Type I CRISPR-Cas system from Shewanella putrefaciens. Nucleic Acids Res., 43, 8913–8923.

16. Otte, K., Kühne, N.M., Furrer, A.D., Baena Lozada, L.P., Lutz, V.T., Schilling, T. and Hertel, R. (2020) A CRISPR-Cas9 tool to explore the genetics of Bacillus subtilis phages. Lett. Appl. Microbiol., 71, 588–595.

17. Gleditzsch, D., Müller-Esparza, H., Pausch, P., Sharma, K., Dwarakanath, S., Urlaub, H., Bange, G. and Randau, L. (2016) Modulating the Cascade architecture of a minimal Type I-F CRISPR-Cas system. Nucleic Acids Res., 44, 5872–5882.

18. Gibson, D.G., Young, L., Chuang, R.Y., Venter, J.C., Hutchison, C.A. and Smith, H.O. (2009) Enzymatic assembly of DNA molecules up to several hundred kilobases. Nat. Methods, 6, 343–345.

19. Schilling, T., Dietrich, S., Hoppert, M. and Hertel, R. (2018) A CRISPR-cas9-based toolkit for fast and precise in vivo genetic engineering of Bacillus subtilis phages. Viruses, 10, 241.

20. Stoiber, M.H., Quick, J., Egan, R., Lee, J.E., Celniker, S.E., Neely, R., Loman, N., Pennacchio, L. and Brown, J.B. (2016) De novo Identification of DNA Modifications Enabled by Genome-Guided Nanopore Signal Processing. bioRxiv.

21. Bailey, T.L., Williams, N., Misleh, C. and Li, W.W. (2006) MEME: Discovering and analyzing DNA and protein sequence motifs. Nucleic Acids Res., 34, W369–W373.

22. Crooks, G.E., Hon, G., Chandonia, J.-M. and Brenner, S.E. (2004) WebLogo: a sequence logo generator. Genome Res., 14, 1188–1190.

23. Rachkevych, N., Sybirna, K., Boyko, S., Boretsky, Y. and Sibirny, A. (2014) Improving the efficiency of plasmid transformation in Shewanella oneidensis MR-1 by removing ClaI restriction site. J. Microbiol. Methods, 99, 35–37.

24. Sorek, R., Lawrence, C.M. and Wiedenheft, B. (2013) CRISPR-mediated adaptive immune systems in bacteria and archaea. Annu. Rev. Biochem., 82, 237–266.

25. Koonin, E. V. and Makarova, K.S. (2019) Origins and evolution of CRISPR-Cas systems. Philos. Trans. R. Soc. B Biol. Sci., 374.

26. Nussenzweig, P.M. and Marraffini, L.A. (2020) Molecular Mechanisms of CRISPR-Cas Immunity in Bacteria. Annu. Rev. Genet., 54, 93–120.

27. Jinek, M., Chylinski, K., Fonfara, I., Hauer, M., Doudna, J.A. and Charpentier, E. (2012) A programmable dual-RNA-guided DNA endonuclease in adaptive bacterial immunity. Science (80-.)., 337, 816–821.

28. Gödeke, J., Paul, K., Lassak, J. and Thormann, K.M. (2011) Phage-induced lysis enhances biofilm formation in Shewanella oneidensis MR-1. ISME J., 5, 613–626.

29. McIntyre, A.B.R., Alexander, N., Grigorev, K., Bezdan, D., Sichtig, H., Chiu, C.Y. and Mason, C.E. (2019) Single-molecule sequencing detection of N6-methyladenine in microbial reference materials. Nat. Commun., 10, 1–11.

30. Anton, B.P., Mongodin, E.F., Agrawal, S., Fomenkov, A., Byrd, D.R., Roberts, R.J. and Raleigh, E.A. (2015) Complete genome sequence of ER2796, a DNA methyltransferase-deficient strain of Escherichia coli K-12. PLoS One, 10.

31. Guzman, L.M., Belin, D., Carson, M.J. and Beckwith, J. (1995) Tight regulation, modulation, and high-level expression by vectors containing the arabinose P(BAD) promoter. J. Bacteriol., 177, 4121–4130.

32. Luo, G.Z., Wang, F., Weng, X., Chen, K., Hao, Z., Yu, M., Deng, X., Liu, J. and He, C. (2016) Characterization of eukaryotic DNA N6-methyladenine by a highly sensitive restriction enzyme-assisted sequencing. Nat. Commun., 7, 1–6.

33. Roberts, R.J., Vincze, T., Posfai, J. and Macelis, D. (2015) REBASE-a database for DNA restriction and modification: Enzymes, genes and genomes. Nucleic Acids Res., 43, D298–D299.

34. Lu, S., Wang, J., Chitsaz, F., Derbyshire, M.K., Geer, R.C., Gonzales, N.R., Gwadz, M., Hurwitz, D.I., Marchler, G.H., Song, J.S., et al. (2020) CDD/SPARCLE: The conserved domain database in 2020. Nucleic Acids Res., 48, D265–D268.

35. Mirdita, M., Schütze, K., Moriwaki, Y., Heo, L., Ovchinnikov, S. and Steinegger, M. (2022) ColabFold: making protein folding accessible to all. Nat. Methods, 10.1038/s41592-022-01488-1.

36. Jumper, J., Evans, R., Pritzel, A., Green, T., Figurnov, M., Ronneberger, O., Tunyasuvunakool, K., Bates, R., Žídek, A., Potapenko, A., et al. (2021) Highly accurate protein structure prediction with AlphaFold. Nature, 596, 583–589.

37. Kempen, M. Van, Kim, S.S., Tumescheit, C., Mirdita, M., Gilchrist, C.L.M., Söding, J. and Steinegger, M. (2022) Foldseek: fast and accurate protein structure search. bioRxiv.

38. Holm, L. (2022) Dali server: structural unification of protein families. Nucleic Acids Res., 50, W210–W215.

39. Berman, H.M., Westbrook, J., Feng, Z., Gilliland, G., Bhat, T.N., Weissig, H., Shindyalov, I.N. and Bourne, P.E. (2000) The Protein Data Bank. Nucleic Acids Res., 28, 235–242.

40. Burley, S.K., Berman, H.M., Bhikadiya, C., Bi, C., Chen, L., Di Costanzo, L., Christie, C., Duarte, J.M., Dutta, S., Feng, Z., et al. (2019) Protein Data Bank: The single global archive for 3D macromolecular structure data. Nucleic Acids Res., 47, D520–D528.

41. Vlot, M., Houkes, J., Lochs, S.J.A., Swarts, D.C., Zheng, P., Kunne, T., Mohanraju, P., Anders, C., Jinek, M., Van Der Oost, J., et al. (2018) Bacteriophage DNA glucosylation impairs target DNA binding by type I and II but not by type V CRISPR–Cas effector complexes. Nucleic Acids Res., 46, 873–885.

42. Malone, L.M., Birkholz, N. and Fineran, P.C. (2021) Conquering CRISPR: how phages overcome bacterial adaptive immunity. Curr. Opin. Biotechnol., 68, 30–36.

43. Liu, Y., Dai, L., Dong, J., Chen, C., Zhu, J., Rao, V.B. and Tao, P. (2020) Covalent Modifications of the Bacteriophage Genome Confer a Degree of Resistance to Bacterial CRISPR Systems. J. Virol., 94.

44. Nielsen, T.K., Forero-Junco, L.M., Kot, W., Moineau, S., Hansen, L.H. and Riber, L. (2022) Detection of nucleotide modifications in bacteria and bacteriophages: Strengths and limitations of current technologies and software. In Molecular Ecology. John Wiley and Sons Inc.

45. Bendall, M.L., Luong, K., Wetmore, K.M., Blow, M., Korlach, J., Deutschbauer, A. and Malmstrom, R.R. (2013) Exploring the roles of DNA methylation in the metal-reducing bacterium shewanella oneidensis MR-1. J. Bacteriol., 195, 4966–4974.

46. Lehman, I.R. and Pratt, E.A. (1960) On the structure of the glucosylated hydroxymethylcytosine nucleotides of coliphages T2, T4, and T6. J. Biol. Chem., 235, 3254–3259.

47. Mathews, C.K. and Allen, J.R. (1983) DNA Precursor Biosynthesis. In Bacteriophage T4 Mathews, C.K. Kutter, E.M. (ed) Washington D.C., USA.

48. Kutter, E., Bryan, D., Ray, G., Brewster, E., Blasdel, B. and Guttman, B. (2018) From host to phage metabolism: Hot tales of phage T4’s takeover of E. coli. Viruses, 10.

