## Supplementary material for "Discovery of a pentose as a cytosine nucleobase modification in *Shewanella* phage Thanatos-1 genomic DNA mediating enhanced resistance towards host restriction systems": Supplemenatry Tables 1-4; Supplementary Figures 1-4

Supplementary Table 1: Bacterial strains used in this study

| **Strain** | **Genotype** | **Purpose** | **Reference** |
|---|---|---|---|
| *Escherichia coli* | | | |
| DH5α-λpir | φ80d*lacZ* ΔM15 Δ(*lacZYA-argF*)U169 $recA_1$ *hsdR17 deoR thi-l supE44 gyrA96* $relA_1$/*λpir* | regular cloning strain | Miller & Mekalanos, 1988 ( doi: 10.1128/jb.170.6.2575-2583.1988) |
| WM3064 | *thrB1004 pro thi rpsL hsdS lacZ* ΔM15 RP4-1360 Δ(*araBAD*) 567Δ*dapA* 1341::[*erm pir*(wt)] | conjugation strain | W. Metcalf, University of Illinois, Urbana-Champaign |
| ER3413 | *fhuA2::IS2, glnX44(AS), λ-, e14-, trp-31, dcm-6, yedZ3069::Tn10, hisG1, argG6, yhdJ11, rpsL104, Δdam-16::KanR, xyl-7, mtlA2, metB1, Δ(mcrC-mrr)114:IS10* | expression strain | E.A. Raleigh, Complete Genome Sequence of ER2796, a DNA Methyltransferase-Deficient Strain of Escherichia coli K-12. PLoS ONE 10(5):e0127446-. |
| BL21-Gold(DE3) | *ompT hsdS(rB – mB – ) dcm+ Tetr gal λ(DE3) endA Hte* | expression strain | Agilent |
| *Shewanella oneidensis* MR-1 | | | |
| S79 | *S. oneidensis* MR-1 wild type | | Venkateswaran et al, 1999 (doi: 10.1099/00207713-49-2-705) |
| S1419 (S6593) | MR-1 ΔLambdaSo ΔMuSo2 (deletion of active prophages) | | Gödeke et al, 2011 (doi: 10.1038/ismej.2010.153) |
| S6733 | S1419 + pCASCADE_RBS-Cas1,wtCas3 + pBAD33 (negative control) | | this work |
| S6735 | S1419 + pCASCADE_RBS-Cas1,wtCas3 + pCRISPR λ SO_2963 | | this work |
| S6737 | S1419 + pCASCADE_RBS-Cas1,wtCas3 + pCRISPR λ SO_2975 | | this work |
| S6741 | S1419 + pCASCADE_RBS-Cas1,wtCas3 + pCRISPR Thanatos TH1_010 | | this work |
| S6743 | S1419 + pCASCADE_RBS-Cas1,wtCas3 + pCRISPR Thanatos TH1_20 | | this work |
| S6086 | S1419 + pTS021 Ara (negative control) | | this work |
| S6197 | S1419 + pTS021 Ara gRNA SO_2963 | | this work |
| S6313 | S1419 + pTS021 Ara gRNA SO_2953 | | this work |
| S6209 | S1419 + pTS021 Ara gRNA TH1_062 | | this work |
| S6210 | S1419 + pTS021 Ara gRNA TH1_126 | | this work |

Supplementary Table 2: Plasmids used in this study

| **Plasmid designation** | **Description** | Reference |
|---|---|---|
| pBBR-MCS5 | broad-range cloning plasmid, GmR | Kovach et al, 1995 (doi: 10.1016/0378-1119(95)00584-1) |
| pBAD33 | pACYC184/p15A; *araC*-$P_{BAD}$; CmR | Guzman et al, 1995 (doi: 10.1128/jb.177.14.4121-4130.1995) |
| pCas1 | Expression vector of the *Shewanella putrefaciens* CN-32 I-Fv type CRISPR-Cas genes *cas7fv*, *cas5fv* and *cas6f* | Gleditzsch et al, 2016 (doi: 10.1093/nar/gkw469 ) |
| pCASCADE_RBS_Cas1 | Cascade genes (*cas7fv cas5fv cas6f*) with a RBS placed in front of *cas7fv* and a FLAG tag added to *cas6f* from pCas1 placed under the control of the ArcA-$P_{BAD}$ promoter cassette from BAD33 in pBBR-MCS5, GmR | this work |
| pCRISPR λ SO_2963 | arabinose-inducible expression of guide RNA targeting LambdaSo gene SO_2963 | this work |
| pCRISPR λ SO_2975 | arabinose-inducible expression of guide RNA targeting LambdaSo gene SO_2975 | this work |
| pTS021 | ori-p15A, *lacPOZ*, *cas9*, $P_{man}$, $P_{van}$, KmR | Otte et al, 2020 (https://doi.org/10.1111/lam.13349) |
| pTS021-ara | ori-p15A, *lacPOZ*, *araC-$P_{BAD}$-cas9*, $P_{man}$, $P_{van}$, KmR | this work |
| pTS021-ara_TH1_062 | gRNA targeting TH1_062 in pTS021-ara | this work |
| pTS021-ara_TH1_126 | gRNA targeting TH1_126 in pTS021-ara | this work |
| pBAD24-TH126 | Arabinose-inducible expression of phage protein TH1_126 | this work |
| pBAD24-NHis-TH1-060 | Arabinose-inducible expression of phage protein TH1_060 | this work |
| pBAD24-NHis-TH1-063 | Arabinose-inducible expression of phage protein TH1_063 | this work |

Supplementary Table 3: Oligonucleotides used in this study

| **Designation/Purpose** | **Sequence (5'→3')** |
|---|---|
| Construction of pCASCADE | |
| pBAD33-AraC-FW | GTGGATCCCCCGGGCTGCAGGTTATGACAACTTGACGGCTAC |
| pBAD33-Pbad-OL-Cas1-RV | GCTAAAATCATCCATGCTAGCCCAAAAAAACGGGTATGG |
| RBS-3xFLAG-FW | CGGGGATCCTCTAGAGAGGAGGTGCATCATGGATTATAAAGATCATGATGGCGATTATAAAGATCATGATATTGATTATAAAGATGATGATGATAAATAAGGCATGCAAGCTTGGC |
| 3xFLAG-RV | GCCAAGCTTGCATGCCTTATTTATCATCATCATCTTTATAATCAATATCATGATCTTTATAATCGCCATCATGATCTTTATAATCCATGATGCACCTCCTCTCTAGAGGATCCCCG |
| pBBR1-MCS5-check-FW | GCGCGTAATACGACTCACTATAGG |
| pBBR1-MCS5-check-RV | CCTCACTAAAGGGAACAAAAGCTGG |
| Cloning of targeting spacers into pBAD33 | |
| λSO_2963-Spacer-FW | CGGGGATCCTCTAGAGGTTCACCGCCGCACAGGCGGCTTAGAAAAGAGGCATTTGTTAAGGGTAACTTTACTTTTGGTTCACCGCCGCACAGGCGGCTTAGAAAGGCATGCAAGCTTGGC |
| λSO_2963-Spacer-RV | GCCAAGCTTGCATGCCTTTCTAAGCCGCCTGTGCGGCGGTGAACCAAAAGTAAAGTTACCCTTAACAAATGCCTCTTTTCTAAGCCGCCTGTGCGGCGGTGAACCTCTAGAGGATCCCCG |
| pBAD33-check-FW | GCCGTCAATTGTCTGATTCG |
| pBAD33-check-RV | GTTTTATCAGACCGCTTCTGCG |
| λSO_2975-Spacer-FW | CGGGGATCCTCTAGAGGTTCACCGCCGCACAGGCGGCTTAGAAAGTACTAGCGGCGGCAGATTTATCGGCTGGAGCGTTCACCGCCGCACAGGCGGCTTAGAAAGGCATGCAAGCTTGGC |
| λSO_2975-Spacer-RV | GCCAAGCTTGCATGCCTTTCTAAGCCGCCTGTGCGGCGGTGAACGCTCCAGCCGATAAATCTGCCGCCGCTAGTACTTTCTAAGCCGCCTGTGCGGCGGTGAACCTCTAGAGGATCCCCG |
| TH_010 (0047)-Spacer-FW | CGGGGATCCTCTAGAGGTTCACCGCCGCACAGGCGGCTTAGAAAACGGTTATAATGCTACTAACATCGCTTCAGGTGTTCACCGCCGCACAGGCGGCTTAGAAAGGCATGCAAGCTTGGC |
| TH_010 (0047)-Spacer-RV | GCCAAGCTTGCATGCCTTTCTAAGCCGCCTGTGCGGCGGTGAACACCTGAAGCGATGTTAGTAGCATTATAACCGTTTTCTAAGCCGCCTGTGCGGCGGTGAACCTCTAGAGGATCCCCG |
| TH_020 (0057)-Spacer-FW | CGGGGATCCTCTAGAGGTTCACCGCCGCACAGGCGGCTTAGAAAAAGTTATGAATCTAATGCTATTTTAGTAGAGAGTTCACCGCCGCACAGGCGGCTTAGAAAGGCATGCAAGCTTGGC |
| TH_020 (0057)-Spacer-RV | GCCAAGCTTGCATGCCTTTCTAAGCCGCCTGTGCGGCGGTGAACTCTCTACTAAAATAGCATTAGATTCATAACTTTTTCTAAGCCGCCTGTGCGGCGGTGAACCTCTAGAGGATCCCCG |
| Construction of pTS021-ara and derivatives (Cas9) | |
| TL 149 | CACTTCCCTGTTAAGTGTACTTATGACAACTTGACGGCTA |
| TL 177 | ATGTCATGACATTGGTGTACACAGTAGAGAGTTGCGATAAA |
| TL 176 | CCACCACTGATTTGAGCGTCAG |

| | |
|---|---|
| TL 163 | CGTCAGATTTCGTGATGCTTGTC |
| RH001 | AGCTTAGGCCCAGTCGAAAG |
| RH002 | CAGCTAGGAGGTGACTGAAG |
| RH003 | ACCGAGCGTTCTGAACAAATCC |
| gRNA TH_062 (T_00098)-f | TACGTAAAAAGAATAGTCGTCGTG |
| gRNA TH_062 (T_00098)-r | AAACCACGACGACTATTCTTTTTA |
| gRNA TH_126 (T_00161)-f | TACGCATGTAGAGCATTTCACTAG |
| gRNA TH_126 (T_00161)-r | AAACCTAGTGAAATGCTCTACATG |
| Cloning of inducible expression of TH1_126 into NcoI site of pBAD24 | |
| pBAD_TH1-126_fw | GCTAGCAGGAGGAATTCACATGATTAATTATAAAGATGGTCTAAATGG |
| pBAD_TH1-126_rev | TCTAGAGGATCCCCGGGTACCTAATAGTTAGTTATCAACACTTCTAC |
| Cloning of inducible expression of TH1_060 and TH1_063 into NcoI site of pBAD24. | |
| pBAD_TH1-060_fwd | GGCTAGCAGGAGGAATTCACATGAAAAAAGTTGTTATTCTTGCTTCTGGTATGG |
| pBAD_TH1-060_rev | TCTAGAGGATCCCCGGGTACTCATAGTTTTTCCCATTGGTGGTCAATATGAC |
| pBAD_TH1-063_fwd | GGCTAGCAGGAGGAATTCACATGAATTTTATCGTCCCTATATTTTCAATGCGT |
| pBAD_TH1_063_rev | TCTAGAGGATCCCCGGGTACTCATGTTACCACCAAATCATTGATTAGAGGC |
| Addition of 6x N-terminal His-tags via primer overhangs | |
| pBAD_split_f_1 | CGTTGCGCAAACTATTAACTG |
| pBAD_NHis_r_1 | GTGGTGATGGTGATGATGCATGTGAATTCCTCCTGCTAGCC |
| pBAD_TH1-60_NHis_f_2 | CATCATCACCATCACCACAAAAAAGTTGTTATTCTTGCTTCTGG |
| pBAD24_split_r_2 | AGTTAATAGTTTGCGCAACG |
| pBAD24_TH1-63_NHis_f_2 | CATCATCACCATCACCACAATTTTATCGTCCCTATATTTTC |

**Supplementary Table 4:** Top 20 structural homology hits from DALI against PDB25 for the predicted protein structure of TH1_063.

| #No | Chain | Z | rmsd | lali | nres | %id | PDB Description |
|---|---|---|---|---|---|---|---|
| 1 | 6kih-A | 18 | 3,4 | 269 | 400 | 8 | TLL1590 PROTEIN; |
| 2 | 4xsu-A | 17,4 | 3,1 | 256 | 368 | 9 | ALR3699 PROTEIN; |
| 3 | 6n1x-A | 17,4 | 3 | 255 | 377 | 8 | GLYCOSYLTRANSFERASE; |
| 4 | 2iv7-A | 15,4 | 3,3 | 249 | 370 | 10 | LIPOPOLYSACCHARIDE CORE BIOSYNTHESIS PROTEIN |
| 5 | 6gng-B | 15,3 | 3,4 | 260 | 526 | 8 | GRANULE-BOUND STARCH SYNTHASE; |
| 6 | 4xyw-A | 15,2 | 3,5 | 241 | 324 | 10 | O-ANTIGEN BIOSYNTHESIS GLYCOSYLTRANSFERASE WBNH; |
| 7 | 2iuy-A | 15,2 | 3,1 | 236 | 340 | 12 | GLYCOSYLTRANSFERASE; |
| 8 | 5zer-A | 14,8 | 3,2 | 244 | 354 | 10 | UDP-GLUCOSE:TETRAHYDROBIOPTERIN GLUCOSYLTRANSFERA |
| 9 | 6eji-A | 14,7 | 3,1 | 239 | 360 | 11 | WLAC PROTEIN; |
| 10 | 2bis-A | 14,6 | 3,8 | 257 | 440 | 12 | GLGA GLYCOGEN SYNTHASE; |
| 11 | 3oka-A | 14,3 | 3,8 | 250 | 378 | 8 | GDP-MANNOSE-DEPENDENT ALPHA-(1-6)-PHOSPHATIDYLINO |
| 12 | 7mi0-A | 14,3 | 3,7 | 244 | 379 | 13 | GLYCOSYLTRANSFERASE; |
| 13 | 4wac-A | 14,2 | 3,6 | 252 | 498 | 11 | GLYCOSYL TRANSFERASE, GROUP 1 FAMILY PROTEIN; |
| 14 | 4pqg-A | 13,8 | 3,2 | 241 | 506 | 10 | GLYCOSYLTRANSFERASE GTF1; |
| 15 | 4n9w-A | 13,8 | 3,3 | 225 | 360 | 8 | GDP-MANNOSE-DEPENDENT ALPHA-(1-2)-PHOSPHATIDYLINO |
| 16 | 5uof-A | 13,8 | 3,7 | 252 | 471 | 10 | ALPHA,ALPHA-TREHALOSE-PHOSPHATE SYNTHASE (UDP-FOR |
| 17 | 5tmb-A | 13,4 | 4,2 | 250 | 451 | 8 | GLYCOSYLTRANSFERASE, OS79; |
| 18 | 4fkz-A | 13,4 | 4,3 | 258 | 384 | 10 | UDP-N-ACETYLGLUCOSAMINE 2-EPIMERASE; |
| 19 | 1xv5-A | 13,2 | 3,5 | 239 | 401 | 8 | DNA ALPHA-GLUCOSYLTRANSFERASE; |
| 20 | 2hy7-A | 13,2 | 4,2 | 238 | 373 | 5 | GLUCURONOSYLTRANSFERASE GUMK; |

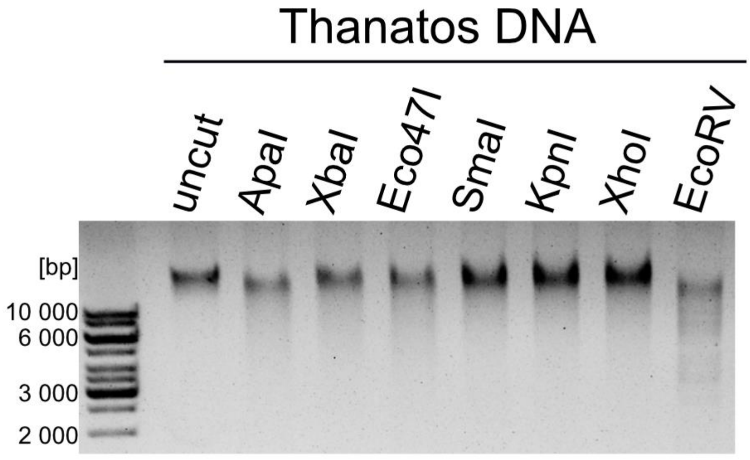


**Supplementary Figure 1: Restriction of *Shewanella* phage Thanatos-1 DNA by type II restriction enzymes.** Shown is a separation of DNA on an 0.8 % (w/v) agarose gel after following 3h of digestion at the appropriate temperatures. As indicated, the following enzymes were used: ApaI (GGGCCC; 12 predicted sites); XbaI (TCTAGA, 75 pred. sites); Eco47I (GGWCC; 96 predicted sites); SmaI (CCCGGG; 11 sites); KpnI (GGTACC; 8 sites); XhoI (CTCGAG; 4 sites); EcoRV (GATATC; 37 sites). Pronounced digestion of Thanatos-1 DNA only occurred when EcoRV was used.

**Characterization of an adenosine 6mA methyltransferase specifically active on NA*TC sequence motifs:** To find out whether base composition upstream or downstream of the ATC trimer influences methylation frequency, we extracted all k-mers containing ATC and extended these both upstream and downstream with up to three occurrences of each base. All instances of the resulting set of k-mers of length 4-6 bp in the ER3413 genome sequence were analyzed with regard to modification frequency. It turned out that methylation signals in the overexpression strain at GATC sites are the strongest, with AATC, TATC and CATC following in decreasing order (Figure 4A). The same analysis conducted for the negative control shows that, firstly, there is no substantial methylation at NATC sites, and that, secondly, there seems to be a sequence context-based effect, with GATC and AATC sites possessing a higher perceived modification rate compared to the other tetramers (Figure 4B). Conclusively, it is highly likely that the higher modification rate at GATC and AATC sites in the overexpression strain results from a context-based bias of tombo's 6mA model, and in reality, there is no preference of any base upstream of the 5'-ATC-3' motif.

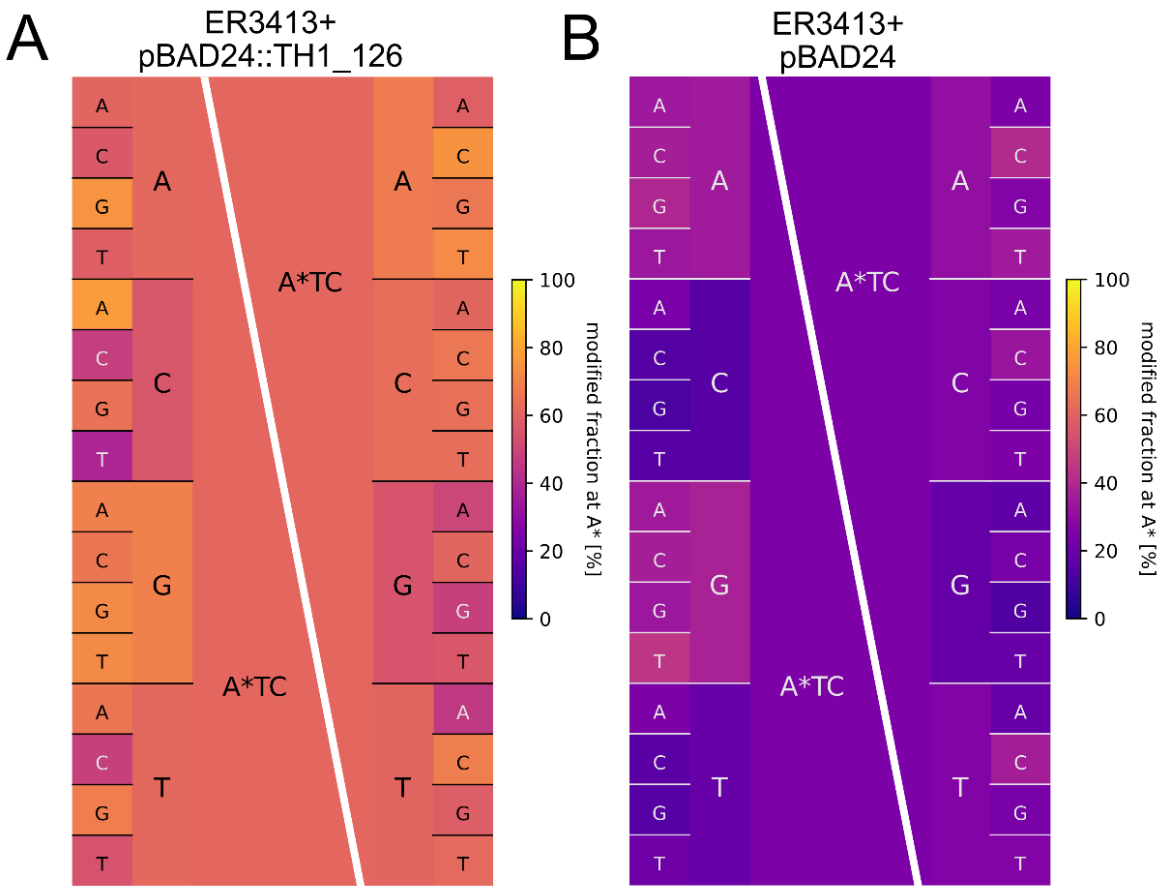


**Supplementary Figure 2: Analysis of all ATC-containing sequences in the ER3413 sequence data with regard to the modified fraction of adenine in the ATC motif.** Comparison of negative control (B) and methylase overexpression strain (A) shows little unspecific predicted modification in the negative control and a substantial predicted fraction of modified bases in the overexpression strain. The ATC core motif is extended upstream and downstream individually by all four bases and the color scale depicts the mean fraction of modified adenines in the respective k-mers as calculated by tombo v1.5 (20).

The *E. coli* MG1655 gDNA with a wild-type methylation profile, which includes dam methylation, was readily cleaved by DpnI as visible by the decrease of DNA fragment lengths, whereas untreated DNA retained its high molecular weight. Integration of empty pBAD24 into ER3413 did not have any effect on the cleavage efficiency of its gDNA. The heterologous expression of TH1_126 via arabinose induction in ER3413, however, led to a shift of the DNA smear towards smaller fragments, providing proof for cleavage of ER3413 gDNA, thereby showing the activity of TH1_126 on E. coli DNA in vivo methylating NATC motifs. Interestingly, although 5'-ATC-3' is part of the EcoRV recognition site (GATATC), the restriction assays of Thanatos-1 DNA showed substantial cleavage by EcoRV (Supplementary Figure 1).

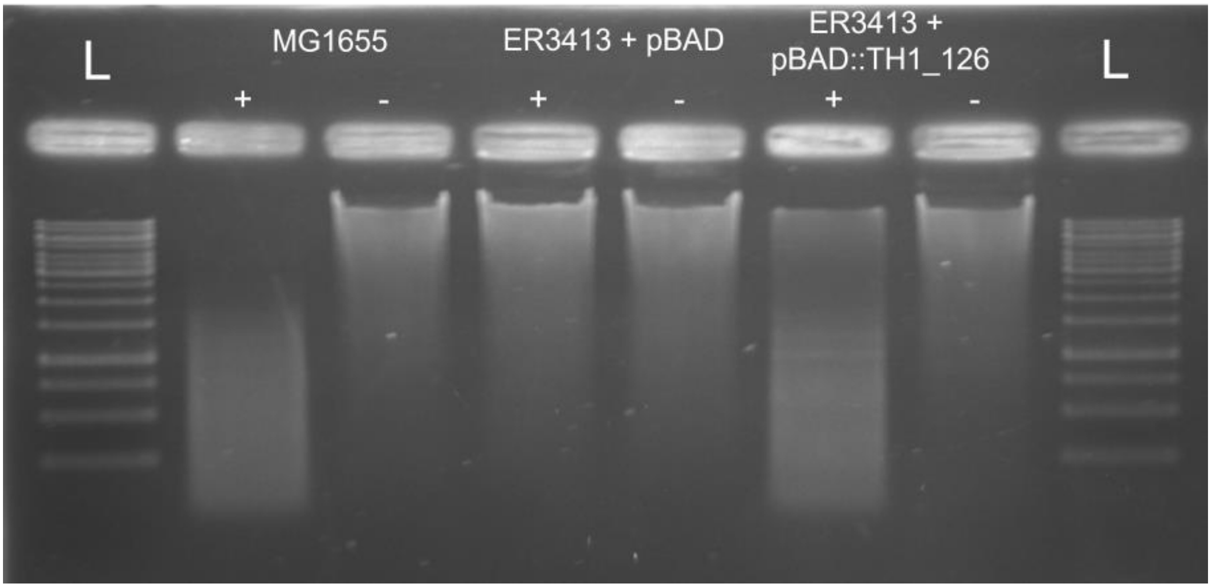


**Supplementary Figure 3: *E. coli* MG1655 gDNA with a wild-type methylation profile, which includes dam methylation, was readily cleaved by DpnI.** This is visible by the decrease of DNA fragment lengths, whereas untreated DNA retained its high molecular weight. Integration of empty pBAD24 into ER3413 did not have any effect on the cleavage efficiency of its gDNA. The heterologous expression of TH1_126 via arabinose induction in ER3413, however, led to a shift of the DNA smear towards smaller fragments, providing proof for cleavage of ER3413 gDNA, thereby showing the activity of TH1_126 on *E. coli* DNA *in vivo* methylating NATC motifs. Interestingly, although 5'-ATC-3' is part of the EcoRV recognition site (GATATC), the restriction assays of Thanatos-1 DNA showed substantial cleavage by EcoRV (Supplementary Figure 1).

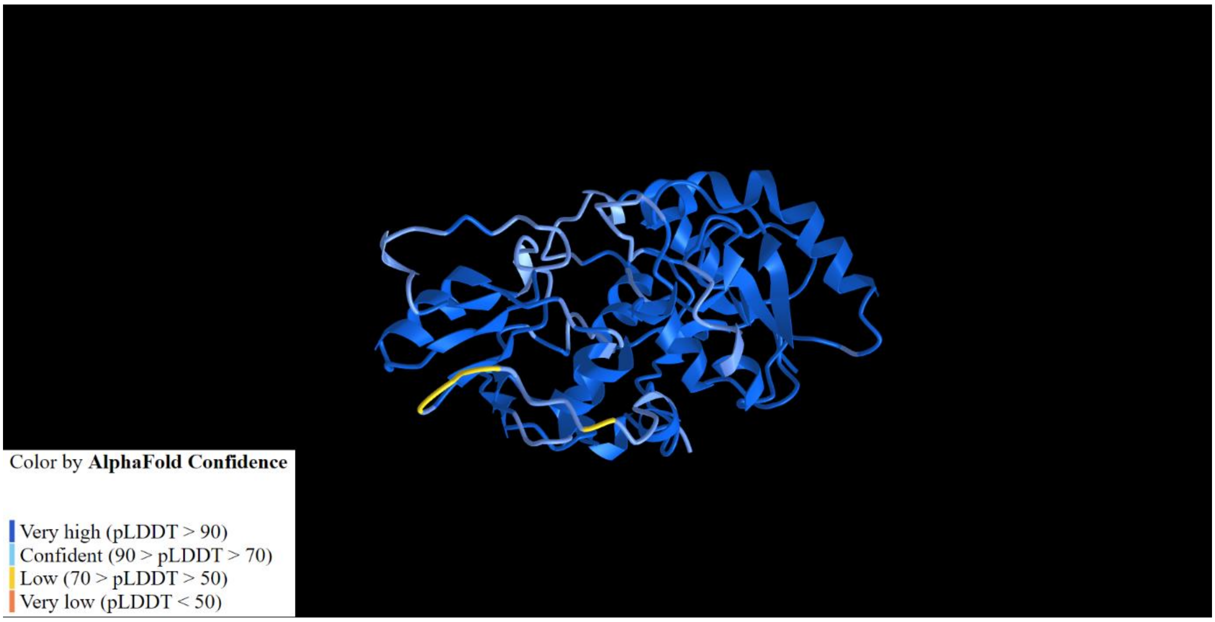


**Supplementary Figure 4: Predicted protein structure of TH1_063 as determined by the ColabFold implementation of the AlphaFold algorithm.** ColabFold outputs five structures ranked by the confidence emitted from the AlphaFold algorithm. This figure shows the top-ranked structure that was also used for DALI search.
